# Rice *NIN-LIKE PROTEIN 1* Rapidly Responds to Nitrogen Deficiency and Improves Yield and Nitrogen Use Efficiency

**DOI:** 10.1101/2020.02.22.961193

**Authors:** Alamin Alfatih, Jie Wu, Zi-Sheng Zhang, Jing-Qiu Xia, Sami Ullah Jan, Lin-Hui Yu, Cheng-Bin Xiang

## Abstract

Nitrogen (N) is indispensable for crop growth and yield, but excessive agricultural application of nitrogenous fertilizers has generated severe environmental problems. A desirable and economical solution to cope with these issues is to improve crop nitrogen use efficiency (NUE). Plant NUE has been a focal point of intensive research worldwide, yet much more has to be learned about its genetic determinants and regulation. Here, we show that rice *NIN-LIKE PROTEIN 1* (*OsNLP1*) plays a fundamental role in N utilization. OsNLP1 protein localizes in nucleus and its transcript level is rapidly induced by N starvation. Overexpression of *OsNLP1* improves plant growth, grain yield and NUE under different N conditions while knockout of *OsNLP1* impairs grain yield and NUE under N limiting conditions. OsNLP1 regulates nitrate and ammonium utilization by cooperatively orchestrating multiple N uptake and assimilation genes. Chromatin immunoprecipitation and yeast-one-hybrid assays show that OsNLP1 can directly bind to the promoter of these genes to activate their expression. Therefore, our results demonstrate that OsNLP1 is a key regulator of N utilization and represents a potential target for improving NUE and yield in rice.

**One-sentence summary:** OsNLP1 rapidly responds to N availability, enhances N uptake and assimilation, and holds great potential in promoting high yield in rice.

## INTRODUCTION

Nitrogen (N) is an essential macronutrient for plants. It is among the most substantial constituents of macromolecular structures like nucleic acids, amino acids, coenzymes, and hormones (Xu et al., 2012). Inorganic form of N such as nitrate also serves as signaling molecule in development (Crawford, 1995). It is axiomatic that absence or limited availability of N will pose deleterious consequences upon plant normal functions, growth and yield (Giehl and von Wirén, 2014). Nevertheless, approximately more than 120 million tons of nitrogenous fertilizers are used annually but less than half of applied fertilizers is utilized by crops, while the leftover in soil is wasted, leading to severe environmental pollution, climate variations and biodiversity loss(Fang et al., 2006). It is challenging to achieve high yields to meet global food demands with reduced inputs of fertilizers. One of the effective solutions to this dilemma is to improve the N use efficiency (NUE) of crops. It is anticipated that improving the crop NUE by merely 1% can significantly elevate yield as well as can reduce the fertilizer input expenses by 1.1 billion US dollars a year (Kant et al., 2012).

NUE is defined as the total amount of yield in the form of grain or biomass achieved per unit of available N (Hawkesford, 2014). The scope of NUE includes the overall processes involved in N uptake, assimilation and utilization by a plant (Xu et al., 2012). N is taken up by plant through nitrate transporters (NRTs) and ammonia transporters (AMTs) from soil mainly in two inorganic forms, nitrate (NO_3_^-^) and ammonium (NH_4_^+^). The absorbed nitrate is initially converted by nitrate reductase (NR) to nitrite and afterwards nitrite is reduced to ammonium by nitrite reductase (NiR) in plastid and/or chloroplast. Ammonium is assimilated into amino acids through the glutamine synthetase/glutamine-2-oxoglutarate aminotransferase (GS/GOGAT) cycle (Wang et al., 2018). Several genes have been identified that could be manipulated for improving nitrogen assimilation in plants. Majority of these identified genes are nitrate and ammonium transporters and key enzymes of N metabolism pathway (Good et al., 2004). Altered expression of these genes has significant effects upon narrow range of functions, with very few possessing broad range effects like overall phenotype or development (McAllister et al., 2012). However, recent reports shed light on high NUE rice breeding. NUE of *indica* rice varieties is 30%–40% higher than that of *japonica* rice varieties (Zhang and Chu, 2020). Three genes, *NRT1.1B* (Hu et al., 2015) *ABC1-1 REPRESSOR1* (*ARE1*) (Wang et al., 2018), and *NR2* (Gao et al., 2019), contribute to the divergence in NUE between the two rice subspecies, indicating a promising strategies to improve the NUE of *japonica* with *indica* genes. Moreover, overexpression of *OsNRT1.1A* in rice not only greatly improves grain yield and N utilization, but also significantly shortens the maturation time (Wang et al., 2018). Another N transporter in the NRT1/PTR FAMILY (NPF), OsNPF6.1^HapB^, is trans-regulated by transcription factor OsNAC42, conferring high NUE and grain yield (Tang et al., 2019). GRF4 (GROWTH REGULATORY FACTOR 4) -DELLA interaction in rice integrates plant growth and regulation of C and N metabolism (Lin et al., 2018).

NODULE INCEPTION (*NLN*), a homologue of assimilation-specific regulator NIT2 in *Chlamydomonas*, is functionally necessary for nodule formation in the legume plant *Lotus japonicas* (Schauser et al., 1999). *NIN* genes encode DNA binding proteins with a bZIP domain of highly conserved 60 amino acid sequence known as RWP-RK motif sequence (Borisov et al., 2003). Homologous genes known as *NLPs* (*NIN*-LIKE PROTEINS) were identified in legumes (Borisov et al., 2003). Through phylogenetic analyses more NLPs were found in other plant species (Chardin et al., 2014; Schauser et al., 2005). Recent studies have highlighted the emerging roles of NLPs in the N signaling and assimilation, root cap release (Karve et al., 2016), seed germination, nodule formation, N and P interaction (Hu et al., 2019). For examples, AtNLP6 and AtNLP7 have central roles in primary nitrate signaling (Konishi and Yanagisawa, 2013; Marchive et al., 2013), and AtNLP8 is involved in crosstalk of nitrate and abscisic acid signaling in seed germination signaling in seed germination (Yan et al., 2016). ZmNLP6, and ZmNLP8 modulate nitrate signaling and assimilation in maize (Cao et al., 2017). NLP1 of *Medicago truncatula* can physically interact with NIN to regulate nitrate-responsive gene expression and the suppression of nodulation by nitrate (Lin et al., 2018). In addition, several components of NLP signaling cascade in *Arabidopsis* have been identified, such as Ca^2+^- DEPENDENT PROTEIN KINASE10/30/32 (CPK10/30/32) (Liu et al., 2017a), NITRATE REGULATORY GENE 2 (NRG2) (Xu et al., 2016), TEOSINTE BRANCHED1/CYCLOIDEA/PROLIFIRATING CELL FACTOR 1-20 (TCP20) (Guan et al., 2017), and NITRATE-INDUCIBLE GARP-TYPE TRANSCRIPTIONAL REPRESSOR 1 (NIGT1) (Maeda et al., 2018).

Most of the studies of NLPs were carried out in *Arabidopsis* and maize. The functions of NLP genes in rice remain largely unknown. According to phylogenic analysis, there are 6 NLPs in rice, among which *OsNLP1* and *OsNLP4* are the closest members with legume *NINs* (Chardin et al., 2014; Schauser et al., 2005). In a recent study, transcriptome analysis revealed the expression of *OsNLP1* was rapidly induced in rice roots by N starvation (Hsieh et al., 2018). Our present study shows that *OsNLP1* play important roles in N utilization and signaling. The expression of *OsNLP1* is induced by N starvation and depressed by either nitrate or ammonium re-supply. OsNLP1 regulates N uptake and assimilation by regulating multiple key N responsive genes related to N metabolism. Moreover, knockout *OsNLP1* significantly impairs plant growth, grain yield and NUE while overexpression of *OsNLP1* enhances NUE and overall grain yield, indicating a potential application of this gene for crop improvement.

## RESULTS

### Expression pattern and subcellular localization of OsNLP1 in response to N availability

To determine the role of OsNLP1 in plant growth and development, we first examined the expression patterns of *OsNLP1* in different tissues and organs of 4-week-old rice plants by quantitative real-time PCR (qRT-PCR). The results showed *OsNLP1* broadly expressed in root, stem, leaf, sheath, flower, panicle and seed, with much higher expression in leaf (Fig. 1A). The tissue expression pattern of *OsNLP1* was further evaluated using *OsNLP1promoter::GUS* transgenic plant. GUS staining was observed in the root, leaf, and panicle (Fig. 1B-H). In root, GUS staining was preferentially localized in the epidermis cells and the vascular tissues (Fig. 1C).

**Figure 1.**
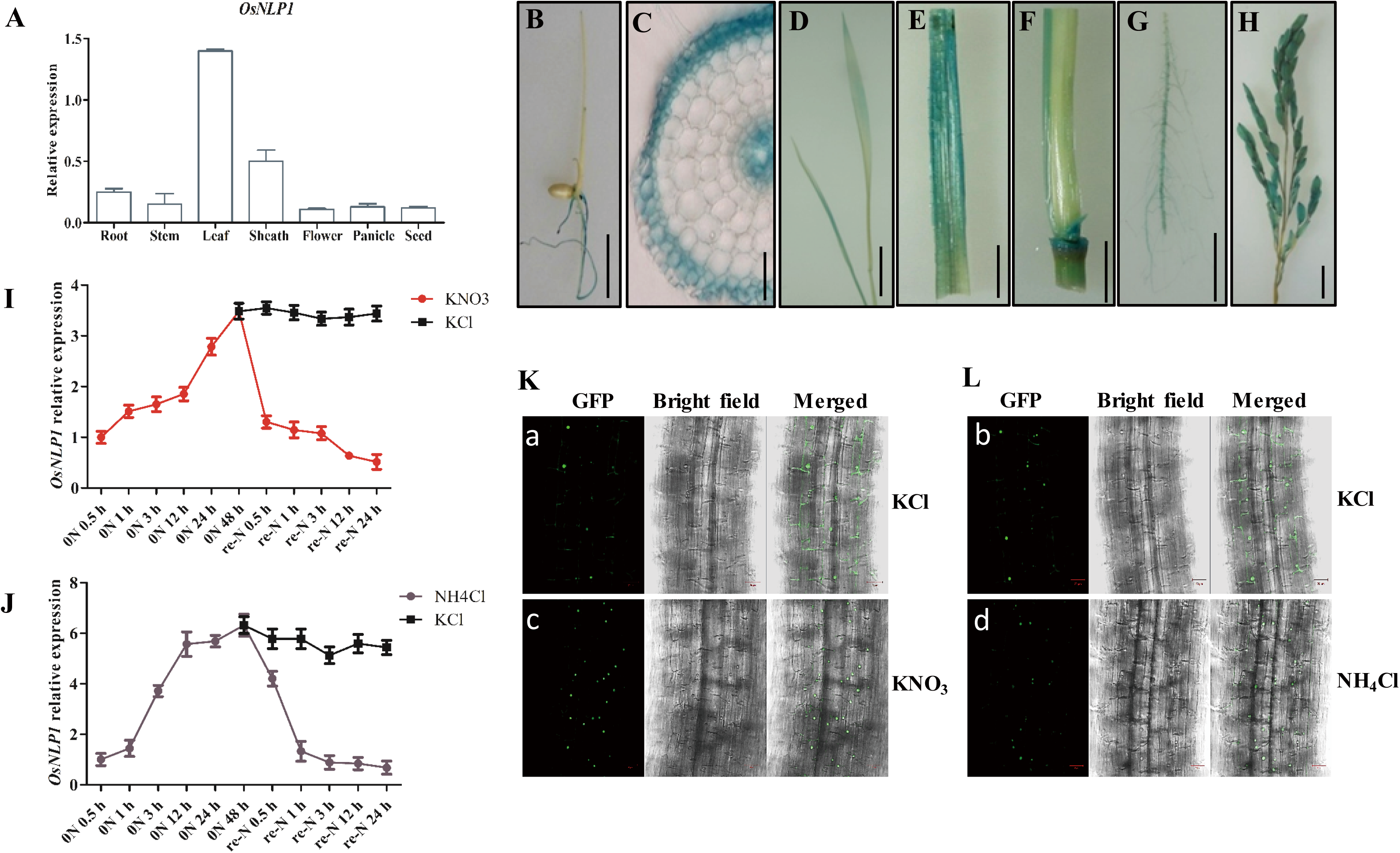
Expression pattern and subcellular localization of OsNLP1 in response to N availability. **A**. Relative expression level of *OsNLP1* in different tissues, as determined using quantitative RT-PCR analysis. *ACTIN* was used as an internal control. Values are the mean ± SD of three replications. **B-H**. GUS staining of *OsNLP1 pro::GUS* transgenic plants. 1-week-old seedling (B, bar = 1 cm). 1-week old seedling root cross section (C, bar = 100 μm). 2-week-old seedling (D, bar = 1 cm). Sheath, stem, root, and panicle of 3-month-old plant (E-H, bar = 1 cm). **I** and **J**. Kinetic analysis of *OsNLP1*expression in response to N starvation and N resupply. Seedling grown on medium contains either 2 mM KNO_3_or 2 mM NH_4_Cl for 1 week, then transferred into medium free of N for 48 hours during this periods seedlings were collected at different time points for RT-qPCR analysis. After 48 hours of N starvation the seedlings were resupplied with either 2 mM KNO_3_ or 2 mM NH_4_Cl or KCl as control for 24 hours, during this time seedlings were collected at different time points for RT-qPCR. *ACTIN* was used as an internal control. Values are the mean ± SD of three replications. **K** and **L**. Localization of OsNLP1 in response to nitrate and ammonium. Confocal laser-scanning microscope observations of NLP1:GFP fusion proteins. Seedling grown on medium free of nitrogen for 2 weeks (a and b). N-starved seedlings were transferred to a nitrate or ammonia medium and observed after 15 minutes (c and d). Scale bars = 20 μm.

To study the responsiveness of *OsNLP1* to N signal, we examined the expression of *OsNLP1* under N starvation. *OsNLP1* transcript level was significantly induced by N starvation. However, when KNO_3_ or NH_4_Cl was resupplied, it was gradually decreased to basal level within 24 hours while KCl control treatment did not alter *OsNLP1* transcript level (Fig. 1I and J), consistent with the previous report that *OsNLP1* was induced by N starvation in rice roots (Hsieh et al., 2018). All these data imply that *OsNLP1* is N responsive and may play some roles in N utilization or signaling

To investigate the subcellular localization of OsNLP1, we generated *35S::OsNLP1-GFP* transgenic lines and found that OsNLP1-GFP was mainly localized in the nucleus with weak signals in cytosol under N starvation (Fig. 1K and L). However, after the addition of either KNO_3_ or NH_4_Cl for 15 minutes, OsNLP1 was significantly induced and mainly localized in the nucleus, with much less in the cytosol (Fig. 1K and L).

### Overexpression of the *OsNLP1* can complement the N starvation phenotype of *Arabidopsis nlp7-1* mutant

As one of homologous genes of *Arabidopsis NLP7*, we wonder whether OsNLP1 plays similar function as AtNLP7. To test this possibility, we overexpressed *OsNLP1* in *Arabidopsis nlp7-1* mutant and found that *OsNLP1* could significantly boost the growth of *nlp7-1* with longer primary roots and increase fresh weight when grown on solid medium containing different N concentrations (1 mM, 3 mM, 10 mM KNO_3_) (Supplemental Fig. S1A-D). The biomass was increased up to 30% in the transgenic plants under 10 mM KNO_3_ condition (Supplemental Fig. S1D). Plant grown in soil also showed that *OsNLP1* can complement the N starvation phenotype of *nlp7-1* (Supplemental Fig. S1E). These results indicate that *OsNLP1* may play as an important N regulator in rice like *NLP7* in *Arabidopsis.*

### OsNLP1 regulates plant growth in response to N availability

To further investigate the functions of *OsNLP1* in rice, we obtained an *OsNLP1* knockdown T-DNA line *osnlp1-1* (*japonica* variety ‘Hwayoung’ background, HY), generated two homozygous loss-of-function mutants (*osnlp1-2* and *osnlp1-3 in* ZH11 background) with CRISPR-Cas9 technology and *OsNLP1*-overexpressing transgenic lines (OE14 and OE16) (Supplemental Fig. S2). All three genotypes were grown in hydroponic culture with three different nitrate concentrations (0.02 mM, 0.2 mM, and 2 mM KNO_3_). A typical seedling growth response is shown in Fig. 2A, indicating that the growth was apparently affected by OsNLP1. The root length was significantly decreased in the *osnlp1-2* and *osnlp1-3* mutants under different nitrate conditions (Fig. 2B). Furthermore, the shoot length of the mutants was significantly reduced under 0.02 mM and 0.2 mM nitrate conditions whereas under high nitrate concentration (2 mM) no obvious difference was seen compared to the WT plants (Fig. 2C). The plant fresh weight of *osnlp1-2* and *osnlp1-3* mutant were significantly decreased under all the different nitrate conditions (Fig. 2D). In contrast, the overexpression lines (OE14 and OE16) exhibited significantly improved root and shoot growth under all nitrate concentrations (Fig. 2B-D).

**Figure 2.**
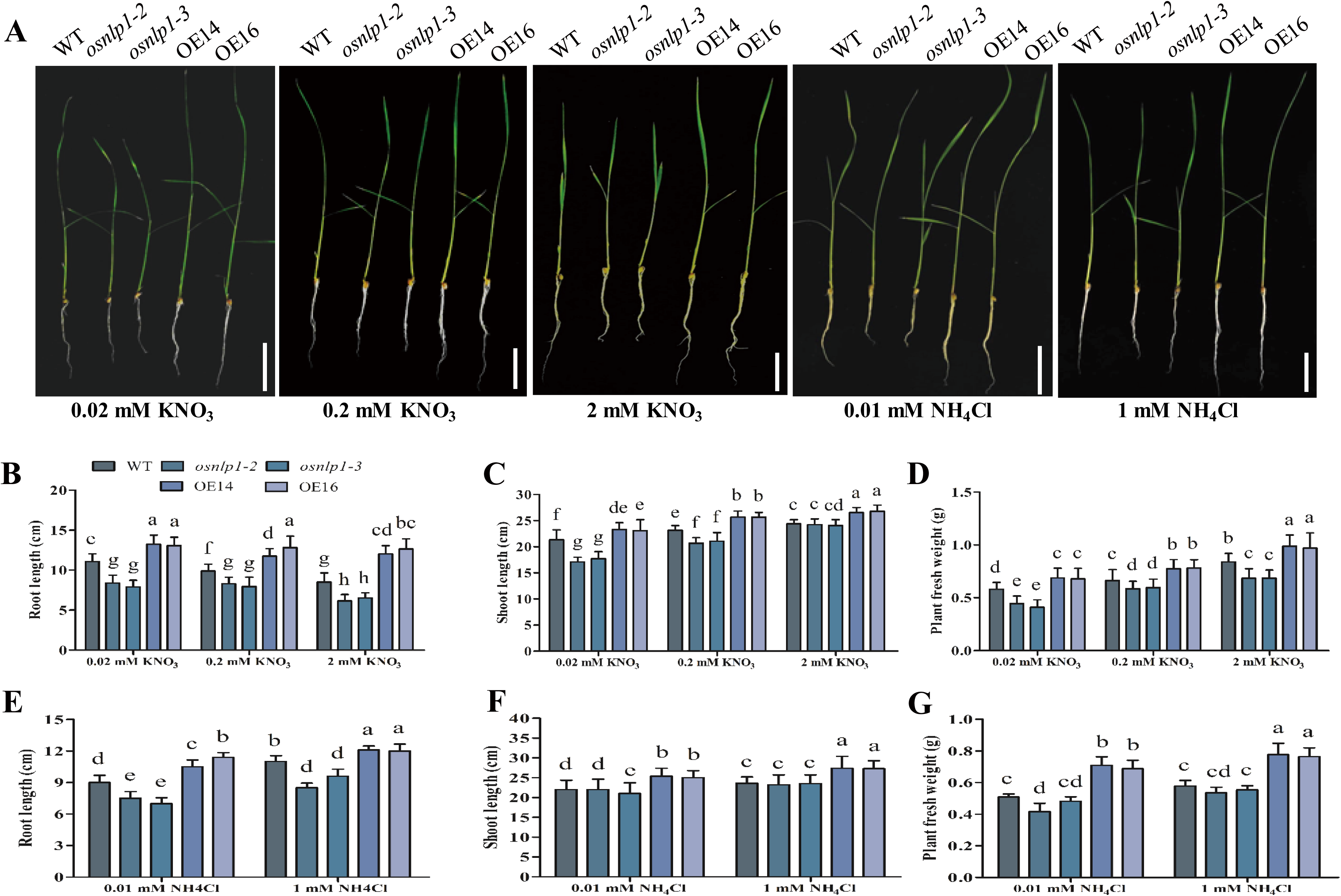
*OsNLP1* improves the growth of plant seedling in hydroponic culture. **A**. Growth of 1-week-old seedlings of wild type (WT), knockout mutants, and overexpression line (OE) at different concentrations of KNO_3_ or NH_4_Cl. The seedlings were hydroponically cultured for 21 days. **B-G**. Measurements of different growth parameters including the shoot length, root length, and total fresh weight. Values are the mean ± SD of three independent replications each containing 16 plants per genotype. Bar = 4 cm. Different letters denote significant differences (P < 0.05) from Duncan’s multiple range tests.

Moreover, we tested the growth response of 3 genotypes to ammonium at 0.01 mM and 1 mM NH_4_Cl (Fig. 2A). Under low and high ammonium concentrations the *osnlp1-2* and *osnlp1-3* mutant showed impaired root growth, while the shoot length showed no difference compared to the WT plants (Fig. 2E and F). The plant fresh weight was significantly decreased in the mutant plants compared to the WT plants (Fig. 2G). In contrast, OE14 and OE16 plants exhibited significantly improved growth under low and high ammonium concentrations with comparison to WT plants as indicated by increased root length, shoot length and plant fresh weight (Fig. 2E-G).

For further confirmation, we checked the T-DNA inserted mutant line *osnlp1-1*. Similarly, seedlings of the *osnlp1-1* mutant exhibited significant growth retardation compared with the wild type HY when grown in hydroponic culture with sufficient or limited N (KNO_3_ or NH_4_Cl) supply (Supplemental Fig. S3). However, under sufficient and limited NH_4_Cl the shoot growth of *osnlp1-1* was significantly impaired unlike *osnlp1-2* and *osnlp1-3* (Supplemental Fig. 3E-G). Taken together, these results demonstrate that OsNLP1 plays crucial roles in N utilization at seedling stage.

### *OsNLP1* is important for grain yield and NUE

To evaluate the performance of *OsNLP1* mutants and overexpression lines in the field, we used three different N concentrations: low (LN), normal (NN), and high N (HN) in the field trial as described in the Methods. A representative plant and panicle were shown in Fig. 3A and B, respectively. Under LN and NN, and HN concentrations, the OE16 line significantly improved NUE by 20.5%, 15%, and 13.6% over the wild type, respectively (Fig. 3C). In contrast, NUE was significantly decreased by about 16% and 14% in *osnlp1-2* mutant under LN and NN concentrations with comparison to wild type plant, respectively (Fig. 3C). Further, the actual yield per plot increased by 21.2% and 11.3% in OE16 under LN and NN, respectively, while it was decreased by 39% and 18.5% in *osnlp1-2* (Fig. 3D). The grain yield per plant showed in Fig. 3E agrees with the actual yield per plot under different N conditions. Moreover, we analyzed several key agronomic traits including tillers number per plant and weight per 1000 seeds. The results show that all these agronomic traits were impaired in *osnlp1-2* while improved in OE16 (Fig. 3F-H). There was no significant difference of plant height between different genotypes except for a slight decrease in plant height of *osnlp1-2* under LN (Fig. 3H). However, under HN condition, all these agronomic traits showed less significant difference among the genotypes (Fig. 3F-H), implying a more important function of OsNLP1 in response to N deficiency than to N rich condition.

**Figure 3.**
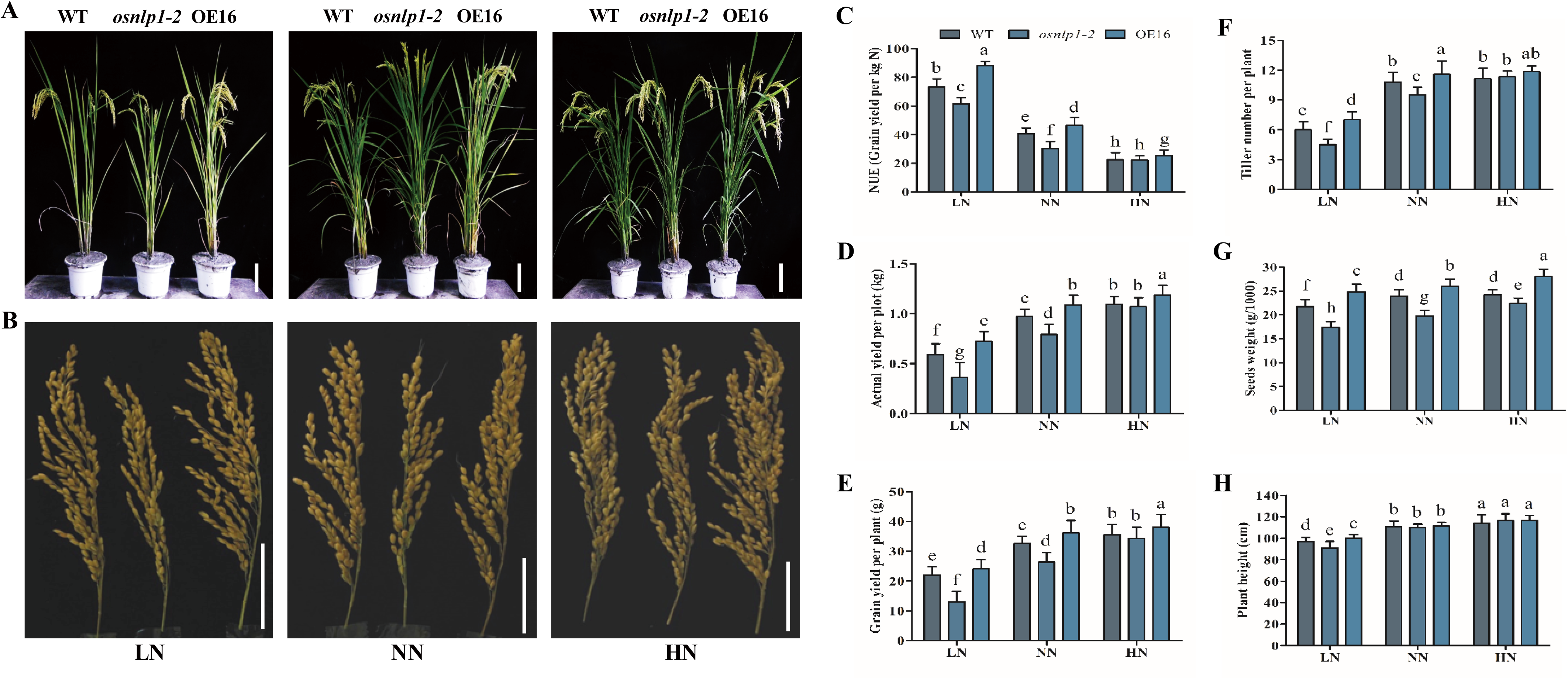
OsNLP1 improves grain yield in the field under different nitrogen levels. **A.** Growth status of a representative WT (ZH11), *osnlp1-2* mutant, and OE16 plant grown in the field of low nitrogen (LN), normal nitrogen (NN), and high nitrogen (HN) conditions. Scale bar =10 cm. **B.** A representative panicle of WT (ZH11), *osnlp1-2*, and OE16 plants. Scale bar =8 cm. **C-H**. Nitrogen use efficiency (NUE), actual yield per plot, grain yield per plant, seeds weight (g/1000), tillers number, and plant height respectively. Values are the means ± SD (30 plants per replicate with 3 replicates). Different letters denote significant differences (P < 0.05) from Duncan’s multiple range tests.

We also did field trial for *osnlp1-1* under normal N condition. The knockdown mutant *osnlp1-1* displayed N deficiency phenotypes with significantly decreases in actual grain yield per plot and grain yield per plant by ∼70% and 66% respectively compared with WT (HY) (Supplemental Fig. S4A-C). Further, *osnlp1-1* mutant shows severe decreases in seed setting rate, weight per 1000 seeds, and plant height (Supplemental Fig. S4D-F). Interestingly, *osnlp1-1* had much more tillers than WT. However, most of those tillers are ineffective tillers (Supplemental Fig. S4A and G). Taken together, these results demonstrate that OsNLP1 plays crucial roles in grain yield and NUE in rice.

### OsNLP1 modulates the expression of N uptake and metabolism related genes

We investigated the role of OsNLP1 in regulating N uptake by using chlorate, a toxic analog of nitrate (Tsay et al., 1993; Wang et al., 1986). Compared with WT, all mutants (*osnlp1-1, osnlp1-2* and *osnlp1-3*) were less sensitive to chlorate with higher survival rates while *OsNLP1*-overexpressing plants were extremely sensitive to chlorate with significantly higher death rates (Fig. 4A and B and Supplemental Fig. S5A and B). Then we found that the acquisition of nitrate and ammonium significantly increased in *OsNLP1-*OE while significantly reduced in *osnlp1* mutants, as demonstrated by ^15^N -nitrate and ^15^N -ammonium feeding experiments (Fig. 4C and D). Consistently, we found that the total N and nitrate contents were reduced in the *osnlp1* mutants while significantly increased in *OsNLP1*-OE plants under both high and low N conditions (Fig. 4E and F, Supplemental Fig. S5C and D). Additionally, NR and NiR activities increased markedly in the *OsNLP1*-overexpressing plants in contrast to the dramatic decrease in the *osnlp1* mutants (Fig. 4G and H, Supplemental Fig. S5E and F). All these results indicate that OsNLP1 is an important regulator of N uptake and assimilation.

**Figure 4.**
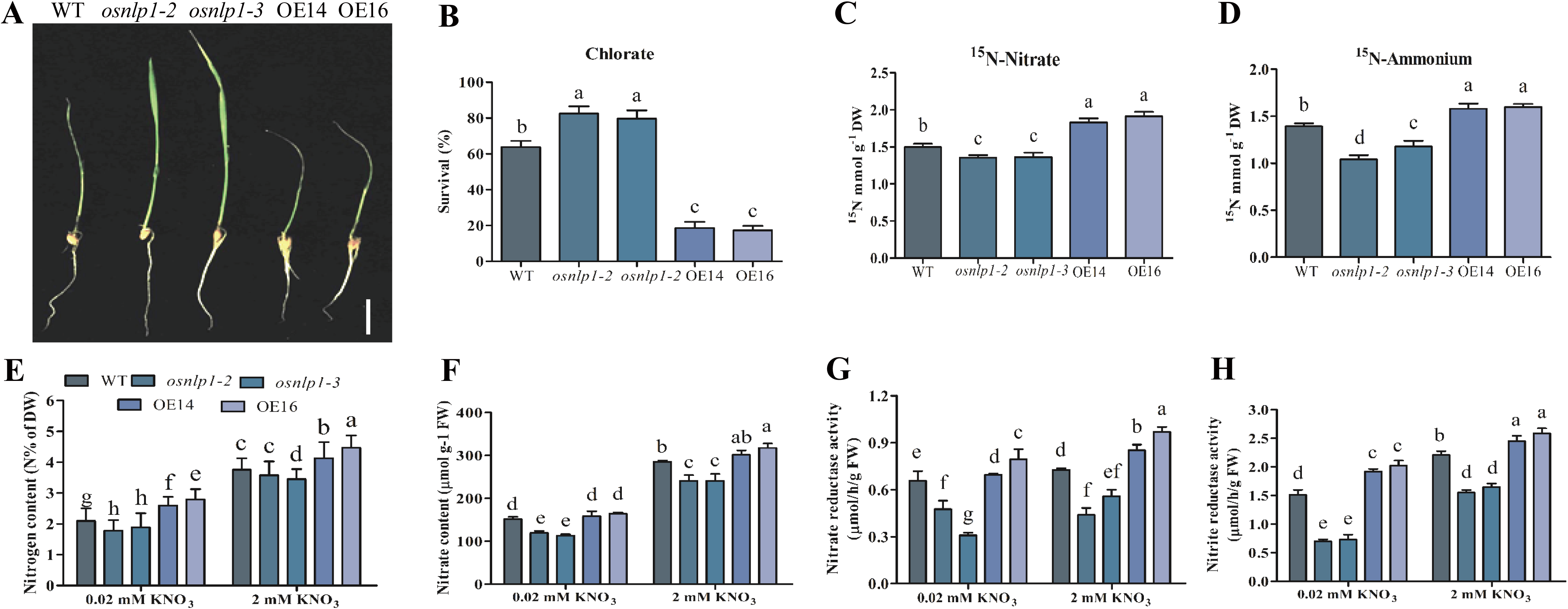
OsNLP1 enhances N uptake and assimilation. **A** and **B**. Chlorate sensitivity and survival rate of 16-day-old plants of WT (ZH11), *osnlp1-2, osnlp1-3*, OE14, and OE16 plants grown on hydroponic culture. Scale bar = 2 cm. Values are the mean ± SD of three independent replications each containing 16 plants per genotype. **C** and **D**. ^15^N accumulation assays in 10 day old seedling of WT (ZH11), *osnlp1-2, osnlp1-3*, OE14, and OE16 plants labeled with ^15^N-nitrate or ^15^N-ammonium. Values are the mean ± SD of three replications. **E-H**. 16-day old plants were grown on hydroponic culture with different N concentrations as described in the Materials and Methods were used for total nitrogen content (E), nitrate content (F), and enzymatic activity (G and H). Values are the mean ± SD of three replications. Different letters denote significant differences (P < 0.05) from Duncan’s multiple range tests. DW, dry weight. FW, fresh weight.

To further assess the role of OsNLP1 in N utilization, we performed qRT-PCR to investigate the expression of key genes related to N uptake and assimilation. Under normal growth condition, the genes responsible for nitrate uptake and transport such as *OsNRT1.1A, OsNRT1.1B, OsNRT2.1*, and *OsNRT2.4*, and for nitrate assimilation such as *OsNIA1, OsNIA2*, and *OsNIA3* were significantly down-regulated in *osnlp1-2, osnlp1-3* mutants but prominently up-regulated in *OsNLP1*-overexpressing plants compared to WT (Fig. 5A-G). An important transcription factor in N signalling, *OsGRF4* was found positively modulated by OsNLP1 (Fig. 5H). Additionally, the expression of genes involved in uptake and assimilation of ammonium, such as *OsAMT1.1, OsAMT2, OsFd-GOGAT, OsGS1.1* and *OsGS2* was similarly regulated by OsNLP1 (Fig. 5I-L). Similar results were revealed in *osnlp1-1* mutant with comparison to the wild type plant HY (Supplemental Fig. S6). Together, these results indicate the importance of OsNLP1 in optimizing utilization of N in response to both nitrate and ammonium

**Figure 5.**
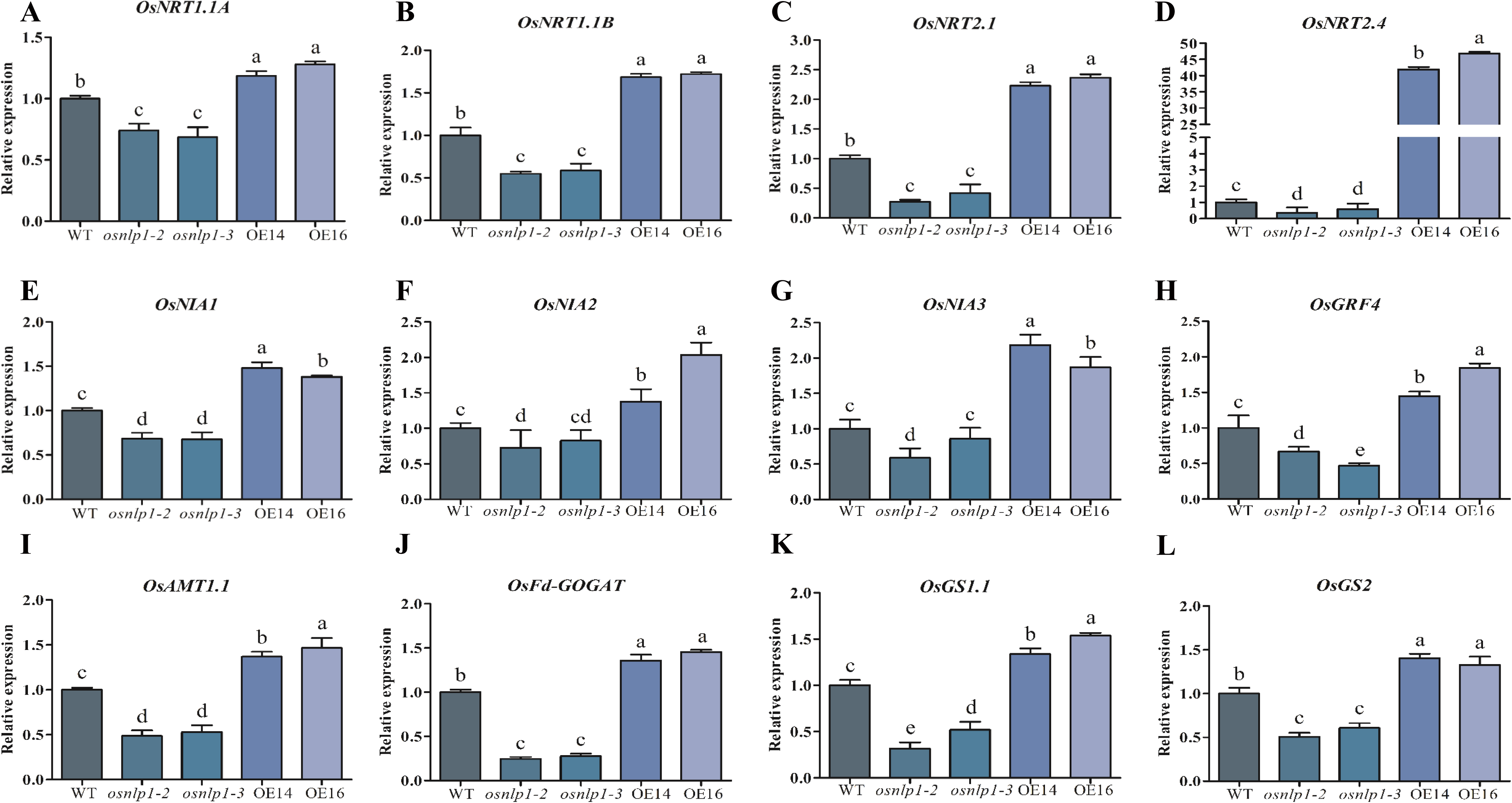
OsNLP1 broadly regulates the genes related to N utilization and signaling. **A-L**. 7-day-old plants grown on medium contains normal N concentration were used for RT-qPCR analysis. *ACTIN* was used as an internal control. Nitrate transporters: *OsNRT1.1A, OsNRT1.1B, OsNRT2.1* and *OsNRT2.4;* nitrate reductase: *OsNIA1, OsNIA2*, and *OsNIA3*; RICE GROWTH FACTOR 4: OsGRF4; ammonium transporter: *OsAMT1.1*; ferredoxin-dependent glutamate synthase: *OsFd-GOGAT*; glutamine synthetases: *OsGS1.1 and OsGS2*. Values are the mean ± SD of three replications. Different letters denote significant differences (P < 0.05) from Duncan’s multiple range tests.

### OsNLP1 directly binds to the nitrate response elements (NREs) in the promoter of N uptake and assimilation related genes

Promoter sequence analysis indicated that there are one or more putative NREs in the promoters of many N responsive genes regulated by OsNLP1 (Supplemental Table S1). Whether these genes are direct targets of OsNLP1 remained to be elucidated. To answer this question, we performed chromatin immunoprecipitation (ChIP) assays using *35S*::*OsNLP1*-*GFP* transgenic plants. ChIP-qPCR showed the promoter fragments containing NRE-like of multiple OsNLP1 regulated N responsive genes, including *OsNRT1.1A, OsNRT1.1B, OsNRT2.4, OsNIA1, OsNIA3, OsAMT1.1* and *OsGRF4*, were significantly enriched in the *35S*::*OsNLP1*-*GFP* transgenic plants compared with the controls (Fig. 6A-G), indicating that OsNLP1 binds to these regions in *vivo*. Furthermore, this result was verified by yeast-one-hybrid assay (Fig. 6H). These data are also consistent with the qRT-PCR data (Fig. 5) and suggest that OsNLP1 directly modulates the expression of these genes.

**Figure 6.**
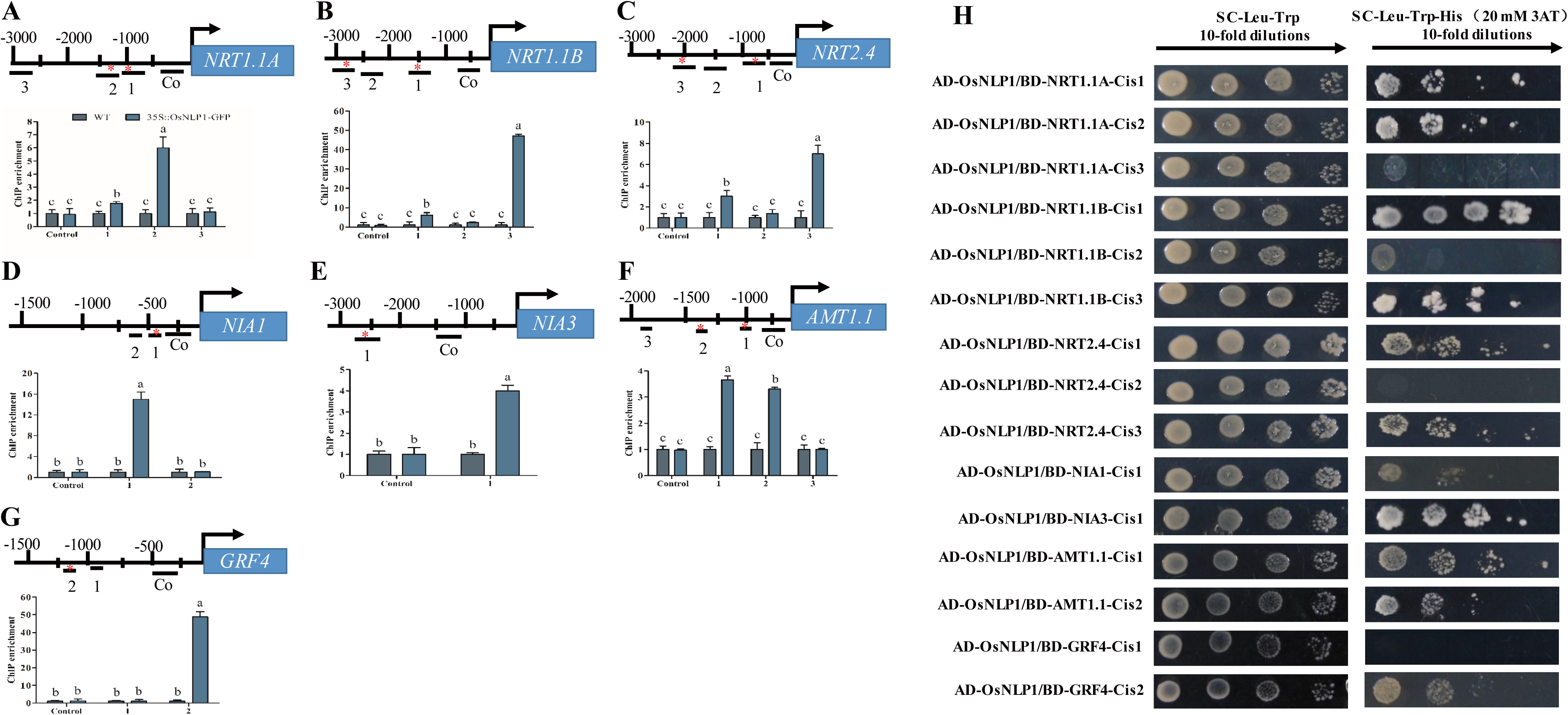
OsNLP1 regulates the expression of multiple N metabolism genes. **A-G**. ChIP–PCR enrichment (relative to wild type) of NRE-like TGACCC…N (9-12)…AAGAG-containing fragments (marked with asterisks) from promoters of *NRT1.1A*(A); *NRT1.1B* (B); *NRT2.4* (C) genes that encode nitrate transporters; *NIA1* (D) and *NIA3* (E) genes that encode NO^−^_3_-assimilation enzymes; *AMT1.1*(F) encodes ammonium transporter; and *GRF4* growth factor (G) in *35S::NLP1-GFP* plants. Values are the mean ± SD of three replications. Different letters denote significant differences (*P* < 0.05) from Duncan’s multiple range tests. **H**. Yeast-one-hybrid assay for OsNLP1 binding to the promoter regions of N metabolism genes and *GRF4*. The pGADT7/NLP1 (AD-NLP1) construct was cotransformed with pHIS2/NIA1/NRT1.1A/NRT1.1B/NIA3/NRT2.4/AMT1.1/GRT4 (BD-cis) separately into yeast strain Y187.

## DISCUSSION

Extensive and intensive researches worldwide have been dedicated to improve NUE in parallel with reduced input of nitrogenous fertilizers. For this purpose, both conventional breeding and genetic engineering have been exploited. The primary focus remained on manipulating individual genes involved in uptake, assimilation and/or metabolism of N (Good and Beatty, 2011). Most of these studies carried out in non-crop plants, with very little showing phenotypic effects on NUE (McAllister et al., 2012). Due to the genetic complexity of NUE, it is urgent to explore new candidate genes that improve NUE and remain stable in crop plants in the field. In this study, we demonstrated that *OsNLP1*, which plays important roles in N utilization, is a promising target gene for crop NUE improvement.

NLP transcription factors have been shown as important regulators in plant growth and development through fine tuning N utilization and signaling pathways (Mu and Luo, 2019). One mechanism of NLPs regulating nitrate-induced gene transcript is mediated by post-translational regulation via a nuclear retention mechanism, such as AtNLP7, ZmNLP6 and ZmNLP8 (Cao et al., 2017; Marchive et al., 2013). Nitrate triggers nitrate-CPK-NLP signaling to phosphorylate AtNLP7, thus enhancing nuclear retention of AtNLP7 (Liu et al., 2017a). However, more and more evidence suggests that, in addition to post-translational regulation, transcriptional regulation of NLPs also plays important roles in the modulation of N signaling and assimilation. On the one hand, many NLPs were showed to respond to the environmental N availability at transcriptional levels. For instance, *AtNLP4, AtNLP5, AtNLP8* and *AtNLP9* in *Arabidopsis* (Chardin et al., 2014), *BnNLP1/4/5/9* subfamilies in *Brassica napus* (Liu et al., 2018), *ZmNLP3* and *ZmNLP7* in maize, (Cao et al., 2017) *TaNLP1* and *TaNLP2* in wheat (Kumar et al., 2018), were found induced by N starvation. *ZmNLP6* and *ZmNP8* in maize, were generally upregulated by N resupply (Cao et al., 2017). In rice, expression of *OsNLP4* is down-regulated by several abiotic stress and induced by low phosphate treatment (Chardin et al., 2014). Recently, microarray analysis found *OsNLP1* was rapidly induced by N starvation in rice root (Hsieh et al., 2018). Consistent to this result, we found that *OsNLP1* is induced by N deficiency while repressed by N resupply (Fig. 1). On the other hand, unlike AtNLP7, ZmNLP6, ZmNLP8, OsNLP3 and OsNLP4, which shuttling between the nucleus and cytoplasm (Cao et al., 2017; Marchive et al., 2013; Wang et al., 2018), AtNLP8 constitutively localizes in nucleus, indicating a different mechanism from that of AtNLP7. Different from AtNLP7, ZmNLP6 and ZmNLP8, which are hardly detected in nucleus under N starvation, we found that OsNLP1 mainly localizes in nucleus with very little in cytosol under N deficient condition. When nitrate or ammonium was resupplied, nucleus localized OsNLP1 rapidly increased with decreased OsNLP1 in cytosol (Fig. 1K and L). Moreover, constitutive expression of *OsNLP1* can significantly activate the expression of N responsive genes as *AtNLP7, ZmNLP6*, and *ZmNLP8* do (Cao et al., 2017; Yu et al., 2016). Taken together, these results suggest that some NLPs respond to N signal at both transcriptional and post-translational levels. OsNLP1 probably plays as a housekeeper to maintain the basal N utilization and at the same time to try to maximize N utilization.

Ammonium rather than nitrate is the major N source for traditionally planted rice in waterlogged fields. However, due to nitrification in the rhizosphere, up to 40% of total N taken up by rice is absorbed as nitrate (Arth et al., 1998; Li et al., 2008). Thus, rice has evolved with the ability to utilize both ammonium and nitrate at high rates. According to the latest studies, NLPs mainly respond to nitrate rather than ammonium, and act as key regulators in nitrate signaling and assimilation. Nucleocytosolic shuttling of AtNLP7 is specific for nitrate rather than ammonium (Marchive et al., 2013), so did ZmNLP6 and ZmNLP8 in maize and *MtNLP1* in *Medicago truncatula* (Cao et al., 2017; Lin et al., 2018). However, our data showed that *OsNLP1* responds to both nitrate and ammonium in a similar manner, induced by nitrate or ammonium starvation while repressed by nitrate or ammonium resupply (Fig. 1). Meanwhile, OsNLP1 positively regulates the transcription of genes related to nitrate uptake and assimilation (*OsNRT1.1A, OsNRT1.1B, OsNRT2.1, OsNRT2.4, OsNIA1, OsNIA2, OsNIA3*) as well as ammonium uptake and assimilation (*OsAMT1.1, OsAMT2, OsFd-GOGAT, OsGS1.1, OsGS2*) and *OsGRF4* as well (Fig. 5). Consistently, *OsNLP1* OE lines have much higher nitrate and ammonium uptake efficiency (Fig. 4A-D) with higher activity of NR and NiR (Fig. 4G-H), whereas *osnlp1* mutants showed the opposite trends (Fig. 4). More importantly, the plant growth was impaired in *osnlp1* mutants while promoted in *OsNLP1* OE plants under conditions with different concentrations of nitrate or ammonium (Fig. 2). All these results demonstrate that OsNLP1 plays a crucial role in both nitrate and ammonium utilization and signaling. As an orchestrator of N response, it is reasonable that OsNLP1 has evolved to respond to both nitrate and ammonium, thus leading to a better resource utilization of rice in the paddy soil.

Genome-wide analyses revealed that AtNLP7 and AtNLP6 modulate a majority of known nitrate assimilation and signalling genes. RWK-RK domains of AtNLPs can directly bind to the NRE motif in the promoter of the N responsive genes to regulate their expression. Similarly, OsNLP1 also orchestrates the transcription of multiple genes related to N uptake such as *OsNRT1.1A, OsNRT1.1B, OsNRT2.1* and *OsAMT1.1*, N assimilation such as *OsNIA1, OsFd-GOGAT, OsGS1.1*, and *OsGS2* (Fig. 5). Further ChIP-qPCR and Y1H results show that OsNLP1 can directly bind to the NREs in the promoter of these genes (Fig. 6). It is worth noting that OsNLP1 can directly regulate *OsNRT1.1A, OsNRT1.1B* and *OsGRF4*, thus indirectly modulates N utilization. All the three genes had been proven as key N utilization regulators important for rice yield and NUE. Overexpression of each of these genes can significantly improve rice yield and NUE in the field (Hu et al., 2015; Li et al., 2018; Wang et al., 2018). Both OsNRT1.1A and OsGRF4 play fundamental roles in maintaining N utilization at high rates for nitrate as well as for ammonium, whereas OsNRT1.1B is responsive for sensing the nitrate signal and triggering nitrate-induced gene expression in the short term (Hu et al., 2015; Li et al., 2018; Wang et al., 2018). Based on these data, it is reasonable to speculate that OsNLP1 can improve grain yield and NUE. Indeed, our field tests confirmed this speculation. Grain yield and NUE were significantly reduced in *osnlp1* mutant under LN and NN conditions while increased in *OsNLP1* OE (Fig. 4). Moreover, *OsNLP1* can functionally complement *Arabidopsis nlp7-1* mutant and improves the plant growth (Supplemental Fig. S1).

In conclusion, our work has demonstrated that OsNLP1 plays a fundamental role in N utilization. *OsNLP1* is responsive to N starvation and orchestrates the expression of nitrate and ammonium uptake and assimilation related genes via the NRE motifs. Overexpression of *OsNLP1* can increase grain yield and NUE in the field, especially under N deficiency condition. Overall, OsNLP1 is a promising candidate for breeding cultivars with high yield and NUE under N-limiting conditions.

## MATERIALS AND METHODS

### Seeds collection and culture conditions

Two *Oryza sativa* L. cultivars: Hwayoung (HY) (Hu et al., 2016) and Zhonghua-11 (ZH11) were used to conduct this study. The T-DNA insertion mutant with HY-background (PFG-2A-50058.R) was obtained from Korea Rice Mutant Center, Pohang, Korea (Jeong et al., 2002). For ZH11 mutant, three independent loss-of-function mutants were generated by CRISPR/Cas9 technique (Hu et al., 2016; Liu et al., 2017b). The overexpression lines (OE14 and OE16) were obtained by amplifying full length cDNA of *OsNLP1* using the primers OsNLP1-OX-F and OsNLP1-OX-R (Supplemental Table S2) and cloned into pCB2006 binary vector. The binary vector was transferred into *Agrobacterium tumefaciens* (EHA105) for rice transformation (Liu et al., 2015). Homozygous lines were selected using glufosinate and expression was confirmed by RT-PCR and qRT-PCR. All the experiments were carried out on ZH11 cultivar or otherwise mentioned.

Plants were grown hydroponically in Kimura B solution (Ehara et al., 1990) and the media was replaced with fresh solution on alternate days. Growth conditions were maintained at 28°C temperature, photo-regime of 16 hours light /8 hours dark, 70% relative humidity, and light intensity at 250 μmol m^-2^ s^-1^.

The *Arabidopsis thaliana* ecotype Columbia (Col-0) was used in this study. *Arabidopsis* seeds were surface-sterilized with 10 % bleach, rinsed five times with sterile distilled water, and placed on half-strength Murashige and Skoog medium containing 1.0% (w/v) sucrose and 0.7% (w/v) agar. After vernalization at 4°C for 2days, seeds were germinated and grown in growth chambers at 22°C and 70% relative humidity. For checking the phenotype in soil, 7-day-old seedlings were transferred to pots containing a 2:1 vermiculite: soil mixture under the condition of an 8-hour light/16-hour dark photoperiod and 120 μmol m^-2^ s^-1^ photon density.

### RNA isolation and qRT-PCR

From the total cellular RNA isolated with Trizol method (Invitrogen, California, USA), 1 μg was subjected for reverse transcription. Step-One Plus Real-Time PCR and TaKaRa SYBR Pre-mix Ex-TaqII reagents kit were employed to compare the expression of *OsNLP1* in WT, mutants, and overexpression lines. The list of primers used in this experiment is shown in Supplemental Table S1.

### Enzyme activity assay

16-day-old hydroponically grown seedlings were used for nitrate reductase (NR) and nitrite reductase (NiR) activity assays. Maximal *in vitro* activity of nitrate reductase was measured according to the method reported earlier (Ferrario-Méry et al., 1998) while enzyme-coupled spectrophotometer assay kit (SKBC, China) was used as per manufacturer’s guidelines to analyze the NiR activity.

### Chlorate sensitivity assay

Seedlings grown in 2 mM KNO_3_ Kimura B solution for four days were treated with 2.0 mM chlorate for 5 days and subsequently recovered in 2.0 mM KNO_3_ provided in Kimura B solution for two days. The survival rate was calculated from control and treated plants.

### ^15^N-nitrate or ^15^N-ammonium uptake and ^15^N accumulation

^15^N-accumulation assay after ^15^N-nitrate or ^15^N-ammonium labeling was performed with ^15^N-labeled KNO_3_ (99 atom % ^15^N, Sigma-Aldrich, no. 335134) or ^15^N-labeled NH_4_Cl (98 atom % ^15^N, Sigma-Aldrich, no. 299251), respectively. For ^15^N-nitrate accumulation assay, rice seedlings were cultured in the Kimura B solution for 10 days. Next, the seedlings were pre-treated with the Kimura B solution for 2 hours and then transferred to modified Kimura B solution containing 5 mM ^15^N-KNO_3_ for 24 hours. At the end of labeling, the roots were washed for 1 minute in 0.1 mM CaSO_4_. Seedlings were collected and dried at 70°C. Finally, the samples were ground and the ^15^N content was measured by continuous-flow isotope ratio mass spectrometer (DELTA V Advantage) with an elemental analyzer (EA-HT, Thermo Fisher Scientific, Inc., Bremen, Germany). For ^15^N-ammonium accumulation assays, the treatment was conducted as above except that 5 mM ^15^N-KNO_3_ was replaced with 1 mM ^15^N -NH_4_Cl and 1 mM KNO_3_. 20 seedlings were collected as one biological replicate, and four biological replicates were used for each treatment.

### GUS activity detection

*OsNLP1*-promotor was cloned in pCB308R (Xiang et al., 1999) The *OsNLP1 pro::GUS* transgenic lines were isolated through glufosinate resistance screening. GUS stained T2 population was produced by following the procedure as described before (Jefferson et al., 1987) and GUS stain solution was prepared according to previously reported procedure (Lei et al., 2007). Stained tissues were de-stained and kept in 70% ethanol. GUS activity in each part was visualized using an Olympus IX81 microscope and a HiROX MX5040RZ digital optical microscope (Questar China Limited).

### Sub-cellular localization

*35S::OsNLP1-GFP* fusion vector was constructed by cloning full length *OsNLP1*-CDS without stop codon in pGWB5 binary vector. The inserted gene was confirmed by gene sequencing followed by transduction of recombinant plasmid into rice callus to achieve transient expression. Green fluorescence of OsNLP1-GFP in transgenic rice root cells was visualized using a ZEISS710 confocal laser scanning microscope: 543 nm for excitation and 620 nm for emission.

### Yeast-one-hybrid assay

The protein putative binding sites and coding sequences were cloned into BD vector (pHIS2) and AD vector (pAD-GAL4-2.1), respectively. The Y1H (yeast-one-hybrid) assay was conducted according to the procedure described earlier (Mao et al., 2016).

### ChIP–qPCR assay

ChIP assay was executed according to the protocol described before (O’Geen et al., 2010) with minor alterations. About 2.0 g of transgenic rice (*35S:OsNLP1-GFP*) seedlings grown in high nitrogen (1.25 mM) for two weeks with provided conditions as 1% formaldehyde (v/v) at 20-25°C in vacuum for 15 minutes was homogenized within liquid nitrogen. Chromatin from lysed nuclei was fragmented ultrasonically to achieve an average length of 500 bp fragments. The anti-GFP antibodies (Sigma, F1804) were immunoprecipitated overnight at 4°C. The immuno-precipitated DNA fragments dissolved in water and kept at -80°C prior to use. The precipitated fragments were used as template for qPCR.

### Field trial of rice

For the field test of *osnlp1-2* mutant and *OsNLP1*-OE (all with ZH11 background), T3 generation plants were grown in Chengdu, Sichuan in 2019 (April to September). The plants density was 10 rows × 20 plants per row for each plot, and four replicates were used for each N condition. Urea was used as the N fertilizer at 80 kg N/hm^2^ for low N (LN), 200 kg N/hm^2^ for normal N (NN), and 500 kg N/hm^2^ for high N (HN). For the field test of *osnlp1-1* in Lingshui, Hainan Island (December 2018 to April 2019), the plants were transplanted in 8 rows ×10 plants per row for each plot with four replicates under normal N condition (100 kg urea/hm^2^). To reduce the variability in field test, the fertilizers were used evenly to every plot for N application level. The plants in the edge were eliminated in each plot in order to avoid margin effects.

### Agronomic traits analyses

Individual plant height, seed setting ratio, tiller number, number of seeds per panicle, and grain yield were measured according to protocol documented earlier (Hu et al., 2015).

### Accession Numbers

Sequence data from this article can be found in the Arabidopsis TAIR database (https://www.arabidopsis.org) or Rice Genome Annotation Project (https://rice.plantbiology.msu.edu/) under the following accession numbers: *AtNLP7, AT4G24020*; *OsNRT1.1A*; *OsNLP1, LOC_*Os03g03900; *LOC_Os08g05910*; *OsNRT1.1B, LOC_Os10g40600*; *OsNRT2.1, LOC_Os02g02170*; *OsNIA1, LOC_Os08g36480*; *OsNIA3, LOC_Os02g53130*; *OsAMT1.1, LOC_Os04g43070*; *OsGOGAT1, LOC_Os01g48960*; *OsGOGAT2, LOC_Os05g48200*; *OsGS1.1, LOC_Os02g50240*; *OsGS2,LOC_Os04g56400.1;* and *OsGRF4, LOC_Os02g47280.*

## SUPPLEMENTAL DATA

Supplemental Fig. S1. *OsNLP1* overexpression in Arabidopsis functionally complements *nlp7-1* mutant and improves growth and fresh weight.

Supplemental Fig. S2. Verification of *osnlp1* mutants and OsNLP1 overexpression lines.

Supplemental Fig. S3. Phenotype of *osnlp1-1* mutant under different N conditions.

Supplemental Fig. S4. The *osnlp1-1* mutant exhibits severely reduced yield in the field.

Supplemental Fig. S5. The *osnlp1-1* exhibits significantly reduced N uptake and assimilation

Supplemental Fig. S6. *OsNLP1* broadly regulates the genes related to N utilization and signaling.

Supplemental Table S1. Predicted NRE sequences in the promoter of NLP-regulated genes.

Supplemental Table S2. Primers used in this study.

## AUTHOR CONTRIBUTIONS

C.B.X., J.W. and A.A. designed the experiments. A.A. and J.W. performed most of the experiments and data analyses. Z.S.Z., X.J.Q. and S.U.J. participated in field trials and data analyses. A.A. wrote the manuscript. C.-B.X. and L.-H.Y. supervised the project and revised the manuscript.

## ACKNOWLEDGEMENTS

This work was supported by grants from the National Natural Science Foundation of China (grant no. 31572183), China Postdoctoral Science Foundation (grant no. 2015M580544), Special Fund of China Postdoctoral Science Foundation (2016T90577) and Ministry of Science and Technology of China (grant no. 2018ZX08009-11B, 2016ZX08005-004-003, and 2016ZX08001003). The authors thank the Korea Rice Mutant Center, Pohang, Korea for providing the T-DNA insertion lines used in this study. Alamin Alfatih and Sami Ullah Jan are recipients of CAS-TWAS President’s Fellowship.

