## supplemental info for "Rice *NIN-LIKE PROTEIN 1* Rapidly Responds to Nitrogen Deficiency and Improves Yield and Nitrogen Use Efficiency"

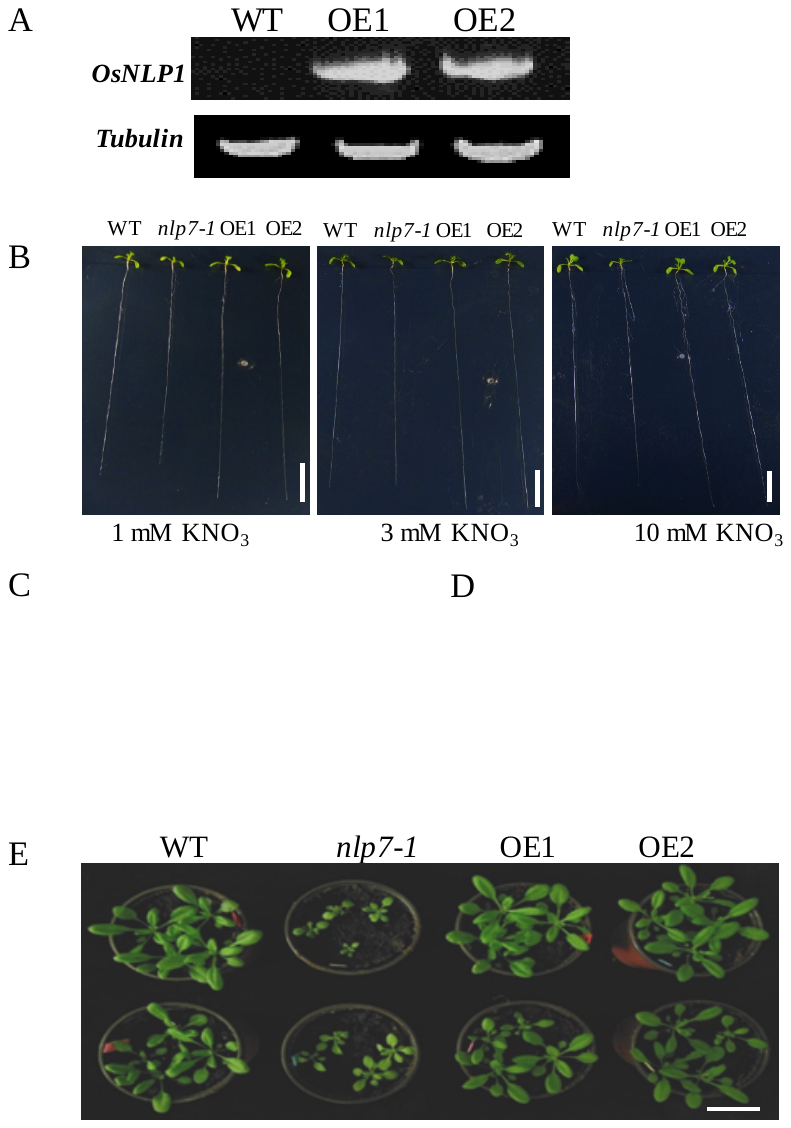

**Figure S1. *OsNLP1* overexpression in Arabidopsis functionally complements *nlp7-1* mutant and improves growth and fresh weight.**

**A**. RNA was extracted from 7-day-old seedlings to perform RT-PCR of transgenic *Arabidopsis* plant expression *35S::OsNLP1. TUBULIN* was used as an internal control**.**

**B**. Growth status of 12-day-old plants of WT, *Arabidopsis nlp7-1* mutant, and transgenic *Arabidopsis* lines overexpressing *OsNLP1* (OE1 and OE2) on a vertical plates containing different nitrogen concentrations. Diameter of the plate is 14.5 cm. Scale bars = 1 cm.

**C** and **D**. Root length and fresh weight of WT, *nlp7-1* mutant, and OE plants. Values are the mean ± SD of five independent replications each containing 6 plants per genotype. Different letters denote significant differences (*P* < 0.05) from Duncan’s multiple range tests.

**E**. WT, *nlp7-1* mutant, and OE plants grown in soil. Scale bars = 8 cm.

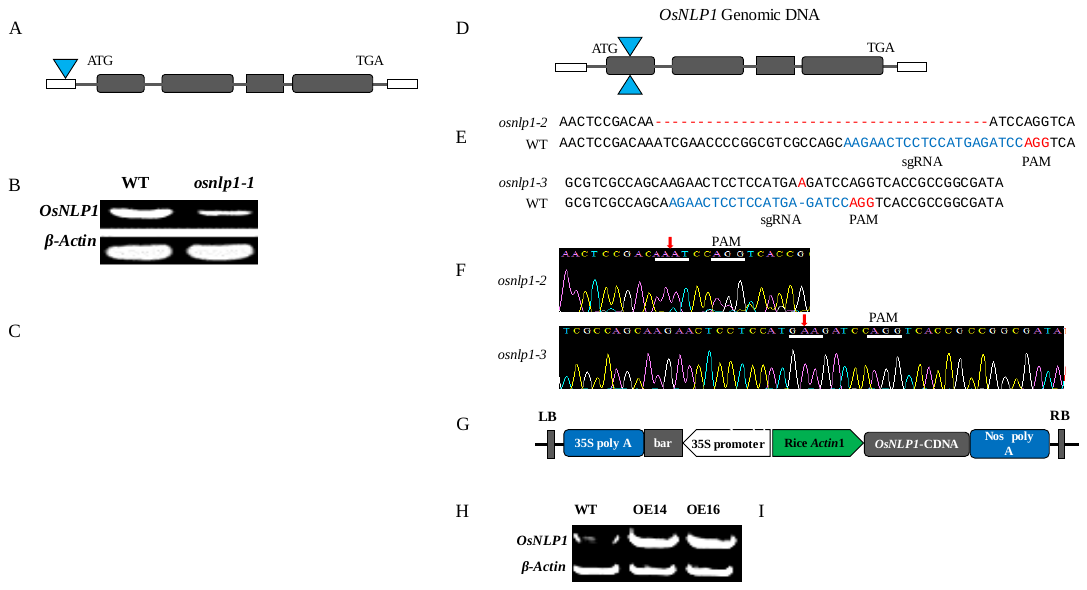

**Figure 2. Verification of *osnlp1* mutants and OsNLP1 overexpression lines.**

**A**. Illustration of the T-DNA insertion site in *OsNLP1*. The triangle indicates the specific location of the T-DNA insertion.

**B** and **C**. Relative expression of the insertion mutant was quantified by semi-quantitative PCR (B), quantitative real-time PCR (C). *ACTIN* was used as an internal reference gene. Values are the mean ± SD of three replications. Different letters denote significant differences (P < 0.05) from Duncan’s multiple range tests.

**D**. Schematic diagram of the CRISPR/Cas9 edited sites in two mutant alleles, named *osnlp1-2* and *osnlp1-3*. Triangles indicate the mutated sites in *OsNLP1*.

**E**. DNA sequences of the edited sites in *OsNLP1*. The dashed line indicates the deletion in the DNA sequences. sgRNA, single guide RNA; PAM, protospacer adjacent motif.

**F**. Sequence confirmation of the edited sites in *osnlp1-2* and *osnlp1-3* mutants. Arrows indicate the edited sites.

**G**. Schematic diagram of the *OsNLP1* overexpression vector.

**H** and **I**. Expression level of *OsNLP1* in overexpression lines OE14 and OE16 in comparison with wild type by semi-quantitative PCR (H), quantitative real-time PCR (I). *ACTIN* was used as internal control. Values are the mean ± SD of three replications. Different letters denote significant differences (P < 0.05) from Duncan’s multiple range tests.

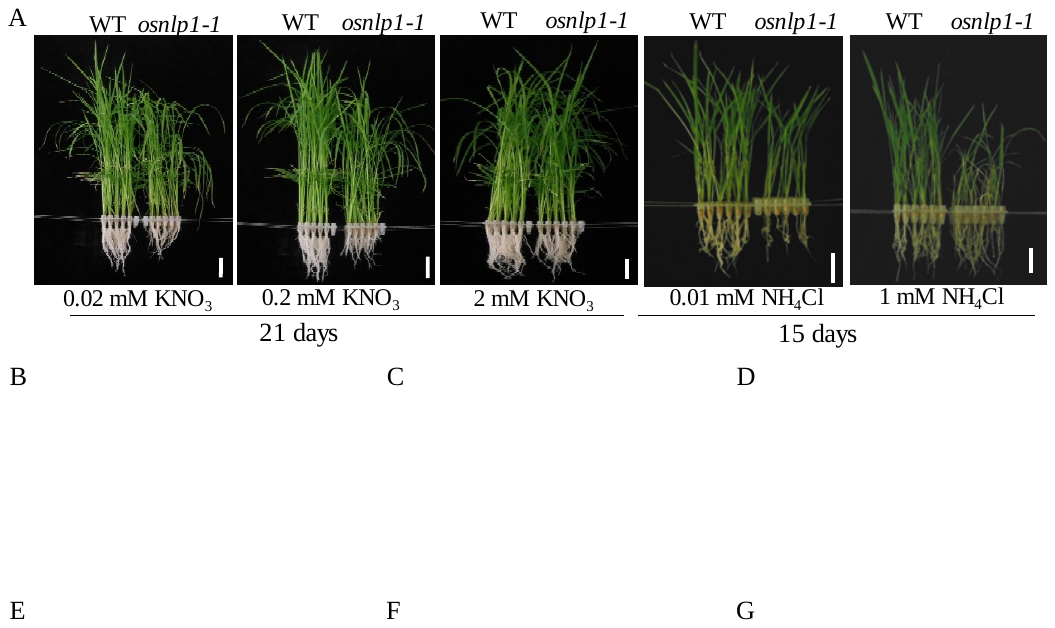

**Figure 3. Phenotype of *osnlp1-1* mutant under different N conditions.**

**A**. Phenotype of WT (HY) and *osnlp1-1* mutant grown in hydroponic medium with different N sources and different concentrations for 21 or 15 days (KNO_3_ or NH_4_Cl). Bar = 4.0 cm.

**B-G**. Measurements of different growth parameters including the shoot length, root length, and total fresh weight. Values are the mean ± SD of three independent replications each containing 16 plants per genotype. Bar = 4 cm. Different letters denote significant differences (P < 0.05) from Duncan’s multiple range tests.

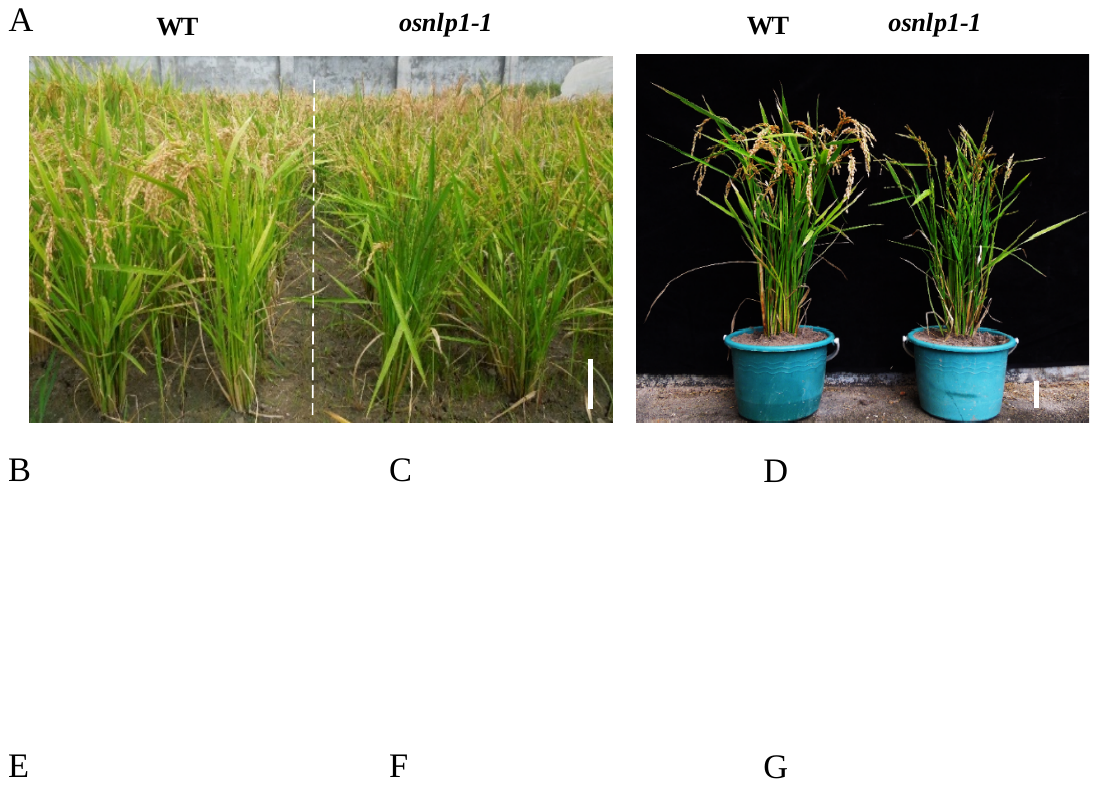

**Figure 4. The *osnlp1-1* mutant exhibits severely reduced yield in the field.**

**A**. Growth status of WT (HY) and *osnlp1-1* mutant plants in the field (Lingshui, Hainan Island, 2019). Bar = 10 cm.

**B-G**. Yield related growth parameters including actual yield per plot, grain yield per single plant, seed setting rate, seeds weight (g/1000), tiller number per plant, and plant height respectively. Values are the means ± SD (30 replicates for plant height and tillers number, 16 replicates for seed setting rate and grain yield per plant, and 3 replicates for actual yield per plot and 1000-grain weight). Different letters denote significant differences (P < 0.05) from Duncan’s multiple range tests.

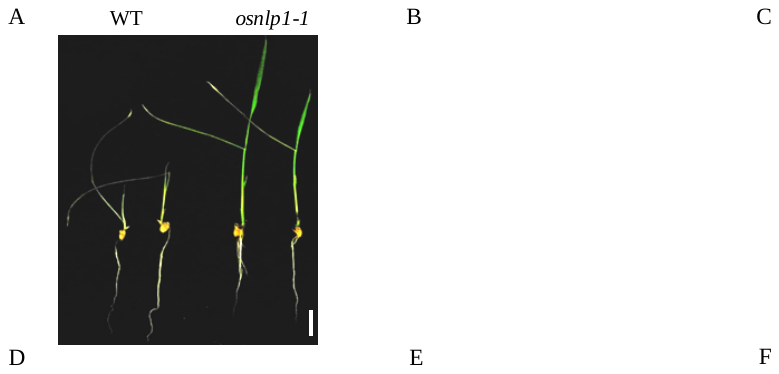

**Figure 5. The *osnlp1-1* exhibits significantly reduced N uptake and assimilation**

16-day-old plants were grown on hydroponic culture with different N concentrations as described in the Materials and Methods were used for chlorate sensitivity, total nitrogen content, nitrate content, and enzymatic activity.

**A** and **B**. Chlorate sensitivity and survival rate of WT (HY) and *osnlp1-1* mutant grown on hydroponic culture. Scale bar = 2 cm. Values are the mean ± SD of three independent replications each containing 16 plants per genotype.

**C** and **D**. Total nitrogen content and nitrate content. Values are the mean ± SD of three replications. Different letters denote significant differences (P < 0.05) from Duncan’s multiple range tests. DW, dry weight. FW, fresh weight.

**E** and **F**. Enzyme activities of nitrate reductase and nitrite reductase in the plants under low and high nitrogen concentrations. Values are the mean ± SD of three replications. Different letters denote significant differences (P < 0.05) from Duncan’s multiple range tests. FW, fresh weight.

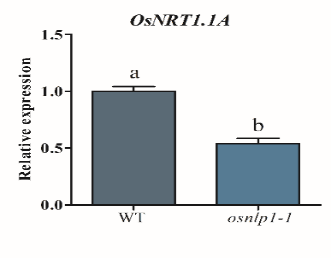

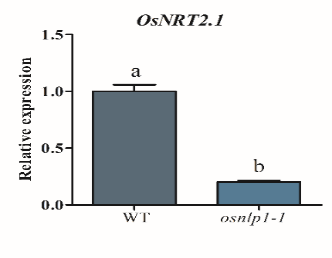

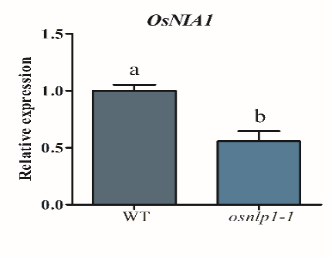

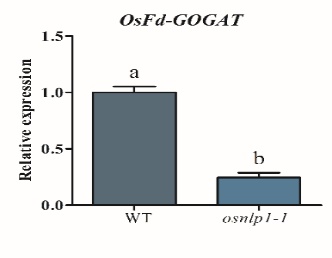

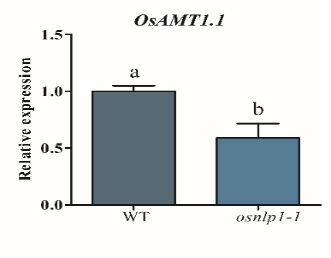

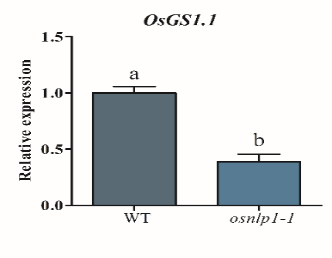

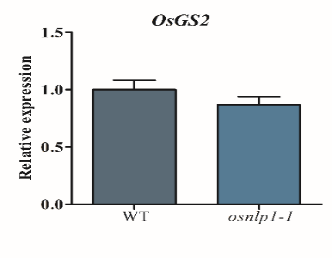

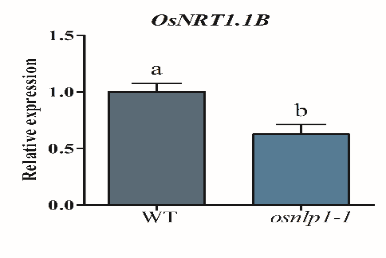

A

B

C

D

E

F

G

H

**Figure 6**. **OsNLP1 broadly regulates the genes related to N utilization and signaling.**

**A-H***.* 7-day-old WT (HY) and *osnlp1-1* plants grown on normal N condition, and then harvested for qRT-PCR analysis. *ACTIN* was used as an internal control. *OsNRT*, nitrate transporter*; OsNIA1*, nitrate reductase1; *OsAMT,* ammonium transporter; *OsFd-GOGAT*, ferredoxin-dependent glutamate synthase;  *OsGS*, glutamine synthetase1.1 respectively. Values are the mean ± SD of three replications. Different letters denote significant Different letters denote significant differences (P < 0.05) from Duncan’s multiple range tests.

**Table S1. Predicted NRE sequences in the promoter of NLP-regulated genes.**

The upper case indicates the conserved nucleotides in NRE motif.

|  | Gene | Gene ID | NRE Sequence |
| --- | --- | --- | --- |
| 1 | *OsNRT1.1A* | *Os08g05910* | NRE*^OsNRT1.1A-1^*: tTGACCCggatcccaact cAcAc  NRE*^OsNRT1.1A-2^*: aTGACtCttgtttagtAcAt  NRE*^OsNRT1.1A-3^*: aTGAtCtctgaccactacAGAcg |
| 2 | *OsNIA1* | *Os08G36480* | NRE*^OsNIA1-1^*: TGgCCttgaattcttgAAGAGc  NRE*^OsNIA1-2^*: TGACgtggcaaaaaaaAAGAG |
| 3 | *OsNIA3* | *Os02G53130* | NRE*^OsNIA3-1^*: TaACCCcagctttaggtgcAAGAaa |
| 4 | *OsGRF4* | *Os02G47280* | NRE*^OsGRF4-1^*: atctTGAaggaaaataaaAAGAG  NRE*^OsGRF4-2^*: tTGACCtacggttgcAtGtcttcg |
| 5 | *OsAMT1.1* | *Os04G43070* | NRE*^OsAMT1.1-1^*: cTGACCttgctgtggcgAActGg  NRE*^OsAMT1.1-2^*: tTGACCagagcaaaaaggttGAGc  NRE*^OsAMT1.1-3^*: tTGAgCCtgattaaacaAAGtct |
| 6 | *OsNRT1.1B* | *Os10G40600* | NRE*^OsNRT1.1B-1^*: cTtACCttgaagggggtAAGAGa  NRE*^OsNRT1.1B-2^*: tTGctCCtttgaatcaAAGAG  NRE*^OsNRT1.1B-3^*: aTGACCCatactgccttAAttac |
| 7 | *OsNRT2.4* | *Os01G36720* | NRE*^OsNRT2.4-1^* cgagCCCtatgtaaaAAaAGgaca  NRE*^OsNR2.4-2^* cgagCCCttgcaaaggttActAGgg  NRE*^OsNRT2.4-3^* cGcCCCaccatctgtcAAatGccc |

**Table S2. Primers used in the study.**

| ***OsNLP1* related primers** |  |
| --- | --- |
| *OsNLP1*-OX-LP | ATGGAGCAGAAGCCGTCGCC |
| *OsNLP1*-OX-RP | TCAAGACAGACCAGTTTGACCG |
| *ACTIN1*-LP | TGGCATCTCTCAGCACATTCC |
| *ACTIN1*-RP | TGCACAATGGATGGGTCAGA |
| *TUBULIN8* LP | CTTAAGCTCACCACTCCAAGCT |
| *TUBULIN8* RP | GCACTTCCACTTCGTCTTCTTC |
| PGWB5-*OsNLP1*-LP | ATGGAGCAGAAGCCGTCGC |
| PGWB5-*OsNLP1*-RP | AGACAGACCAGTTTGACCGAA |
| *OsNLP1*-promoter- GUS-LP | ATTTATACGCTGGATCCGTGG |
| *OsNLP1*-promoter-GUS-RP | GACCGGACAAGCACCACCA |
| **Nitrogen related gene primers** |  |
| *OsNRT1.1A*-qRT-PCR -LP | CCCACACCAAGCAATTCAGG |
| *OsNRT1.1A*- qRT-PCR RP | GTCTTCACCTCCTCCACGTC |
| *OsNRT1.1B*- qRT-PCR LP | GGCAGGCTCGACTACTTCTA |
| *OsNRT1.1B* - qRT-PCR RP | AGGCGCTTCTCCTTGTAGAC |
| *OsNRT2.1*- qRT-PCR LP | ACGGCACAAAGTACAAGACG |
| *OsNRT2.1*- qRT-PCR RP | CCACTGCGGGAAGTAGATG |
| *OsNRT2.4-* qRT-PCR LP | TTCGTCGCGCTCCGGTTCG |
| *OsNRT2.4*- qRT-PCR RP | CGCACGGGAGTAGGTAGGTG |
| *OsNIA1-* qRT-PCR LP | CTGCCTCACCAAGGACAG |
| *OsNIA1*- qRT-PCR RP | TTCCTACTCCTCGTCCTCCT |
| *OsNIA2*- qRT-PCR LP | GAACGAGGAGTAGGAGCACA |
| *OsNIA2*- qRT-PCR RP | GGGCTACAAGATCAAACCAA |
| *OsNIA3*- qRT-PCR LP | GCTCAAGCGCATCATCGTCG |
| *OsNIA3*- qRT-PCR RP | GTAGGCGTATCCCTTCATCGTA |
| *OsGRF4* qRT-PCR LP | ATGACCGCGAGGTGGCCGC |
| *OsGRF4* qRT-PCR RP | CGAAGTACGGACCATATCCAAG |
| *OsAMT1.1*- qRT-PCR LP | GGTTTCTCTCCCTCTCCGAT |
| *OsAMT1.1*- qRT-PCR RP | CCACCTTCACACCACACATT |
| *OsFd-GOGAT* qRT-PCR LP | TGGTCTCCGCCCAGCAC |
| *OsFd-GOGAT* qRT-PCR RP | CAGTTTGTAGGTCAACCGTTATCAT |
| *OsGS1.1*- qRT-PCR LP | ATGGCACCGGTGAAGGTGTTTG |
| *OsGS1.1*- qRT-PCR RP | GTCCTGGAAAGCTGGCATTTGA |
| *OsGS2*- qRT-PCR LP | ATGGCACCGGTGAAGGTGTTTG |
| *OsGS2*- qRT-PCR RP | TCGAAGAGAAGCAGATCCCCG |
| **ChIP related gene primers** |  |
| CHIP-*OsNRT1.1A*- control-LP1 | CCATGGCTATATATACTTCGC |
| CHIP-*OsNRT1.1A*- control-RP1 | CACCACTAATCAATCATCACCAC |
| CHIP-*OsNRT1.1A*-LP1 | TGAACTTGCTCGACTTGGTGT |
| CHIP-*OsNRT1.1A*-RP1 | GAACTTCACCACCGGGTTGT |
| CHIP-*OsNRT1.1A*-LP2 | ACAACCCGGTGGTGAAGTTC |
| CHIP-*OsNRT1.1A*-RP2 | CCTATCCAATTGATCGATGACT |
| CHIP-*OsNRT1.1A*-LP3 | TGCTGCTAATGAGTCAAGTTGG |
| CHIP-*OsNRT1.1A*-RP3 | CTAAGTATCCCGCATGATCTCT |
| CHIP-*OsNRT1.1B*-control-LP1 | AGTGGTACTACGACGAGTGTA |
| CHIP-*OsNRT1.1B*- control-RP1 | GTTGAAATTGAAATGATATGATGG |
| CHIP-*OsNRT1.1B*-LP1 | AATTCCTATCCTTTCAAATCTAATT |
| CHIP-*OsNRT1.1B*-RP1 | CACATTGTATGAGGACTAATGG |
| CHIP-*OsNRT1.1B*-LP2 | CATCTGCATGTCCATTTCTACA |
| CHIP-*OsNRT1.1B*-RP2 | AGAGACTGTGTGGAGCCATG |
| CHIP-*OsNRT1.1B*-LP3 | AGACAGGCAGCCTGCCAGA |
| CHIP-*OsNRT1.1B*-RP3 | CATGCACTGCACGTCATTGC |
| CHIP-*OsNIA1*-control-LP1 | GAAGACAATGGAACGAGTGGT |
| CHIP-*OsNIA1*- control-RP1 | CAAATTGTGTACCCAGTCAATC |
| CHIP-*OsNIA1*-LP1 | AGATTCACACTACTATTATACCAA |
| CHIP-*OsNIA1*-RP1 | TGGCAAGGTGGCTCTGGTG |
| CHIP-*OsNIA1*-LP2 | CACCAGAGCCACCTTGCCA |
| CHIP-*OsNIA1*-RP2 | GGCAAGCAAGGCAGGCAGG |
| CHIP-*OsNIA3*-control-LP1 | GAACTCTATACTCTACTTCTAATA |
| CHIP-*OsNIA3*- control-RP1 | GAGTCTAGAGTCTATACAGGAA |
| CHIP-*OsNIA3*-LP1 | AACACAGGCTGATGGATTAGTT |
| CHIP-*OsNIA3*-RP1 | CAGGCAAGCTGCATTCGGG |
| CHIP-*OsGRF4*-control-LP1 | GCGGCGTGGTACGAGCAG |
| CHIP-*OsGRF4*- control-RP1 | TATCCATCTATCCATCCATTCC |
| CHIP-*OsGRF4*-LP1 | TTAGTCTGCTGCTCCAACATC |
| CHIP-*OsGRF4*-RP1 | TTCAATTTCAACCAAACTTCTAATT |
| CHIP-*OsGRF4*-LP2 | AAGCTGTAAGTTTGAAGGAAAAG |
| CHIP-*OsGRF4*-RP2 | CGATGGCAACAGTGCATGAG |
| CHIP-*OsNRT2.4*- control-LP1 | GTCTATCTCTCTTATCCGATTC |
| CHIP-*OsNRT2.4*-control-RP1 | CCTTAGCCGATTTGGACTGC |
| CHIP-*OsNRT2.4*-LP1 | AGTCGTGTCCATTATGGGCG |
| CHIP-*OsNRT2.4*-RP1 | GCTCAACCCGAGAGCACGA |
| CHIP-*OsNRT2.4*-LP2 | GGCCGATGAGTATCATAACCT |
| CHIP-*OsNRT2.4*-RP2 | AAGTCCTTGGGTACTACTTTCT |
| CHIP-*OsNRT2.4*-LP3 | TAGCATCAGGGAATAAGTCTGT |
| CHIP-*OsNRT2.4*-RP3 | CGGCTATTGAACATCCCTGG |
| CHIP-*OsAMT1.1*-LP1 | CATCCTTGCTGATGGCGATTT |
| CHIP-*OsAMT1.1*-RP1 | TGTGGAGACCCGACGCAGT |
| CHIP-*OsAMT1.1*-LP2 | CCAGAAGAGATGTTACTGTTGA |
| CHIP-*OsAMT1.1*-RP2 | CATCGAAGAGGACTTTGGTGT |
| CHIP-*OsAMT1.1*- control-LP3 | ACACCAAAGTCCTCTTCGATG |
| CHIP-*OsAMT1.1*- control-RP3 | CGCACAGATTGACAAACAGTTT |
| **Yeast-one-hybrid related gene primers** |  |
| PHIS2-*OsNIA1*-LP1 | CTGGCCTTGAATTCTTGAAGAGCA |
| PHIS2-*OsNIA1*-RP1 | CGCGTGCTCTTCAAGAATTCAAGGCCAGAGCT |
| PHIS2-*OsNIA1*-LP2 | CCTCTTTTTTTTTGCCACGTCAA |
| PHIS2-*OsNIA1*-RP2 | CGCGTTGACGTGGCAAAAAAAAAGAGGAGCT |
| PHIS2-*OsGRF4*-LP2 | CCGAAGACATGCAACCGTAGGTCAAA |
| PHIS2-*OsGRF4*-RP2 | CGCGTTTGACCTACGGTTGCATGTCTTCGGAGCT |
| PHIS2-*OsNRT1.1B*-LP2 | CCTCTTTGATTCAAAGGAGCAAA |
| PHIS2-*OsNRT1.1B*-RP2 | CGCGTTTGCTCCTTTGAATCAAAGAGGAGCT |
| PHIS2-*OsNRT1.1B*-LP3 | CATGACCCATACTGCCTTAATTACAA |
| PHIS2-*OsNRT1.1B*-RP3 | CGCGTTGTAATTAAGGCAGTATGGGTCATGAGCT |
| PHIS2-*OsNRT1.1A*-LP1 | CTGACCCCCTCCCAAGGTTTGTAAGA |
| PHIS2-*OsNRT1.1A*-RP1 | CGCGTCTTACAAACCTTGGGAGGGGGTCAGAGCT |
| PHIS2-*OsNRT1.1A*-LP2 | CATGATAATGATGCAAAAAAGAGGA |
| PHIS2-*OsNRT1.1A*-RP2 | CGCGTCCTCTTTTTTGCATCATTATCATGAGCT |
| PHIS2-*OsAMT1.1*-LP2 | CTTGACCAGAGCAAAAAGGTTGAGCA |
| PHIS2-*OsAMT1.1*-RP2 | CGCGTGCTCAACCTTTTTGCTCTGGTCAAGAGCT |
| PHIS2-*OsAMT1.1*-LP3 | CTTGAGCCTGATTAAACAAAGTCTA |
| PHIS2-*OsAMT1.1*- RP3 | CGCGTAGACTTTGTTTAATCAGGCTCAAGAGCT |
| PHIS2-OsNRT2.4-LP1 | CCGAGCCCTATGTAAAAAAAGGACAA |
| PHIS2-OsNRT2.4-RP1 | CGCGTTGTCCTTTTTTTACATAGGGCTCGGAGCT |
| PHIS2-*OsNRT2.4*-LP3 | CCGCCCCACCATCTGTCAAATGCCCA |
| PHIS2-*OsNRT2.4*-RP3 | CGCGTGGGCATTTGACAGATGGTGGGGCGGAGCT |
| PHIS2-*OsNIA3*-LP1 | CTAACCCCAGCTTTAGGTGCAAGAAAA |
| PHIS2-*OsNIA3*-RP1 | CGCGTTTTCTTGCACCTAAAGCTGGGGTTAGAGCT |
